# *HvSL1* and *HvMADS16* promote stamen identity to restrict multiple ovary formation in barley

**DOI:** 10.1101/2022.10.14.512235

**Authors:** Caterina Selva, Xiujuan Yang, Neil J. Shirley, Ryan Whitford, Ute Baumann, Matthew R. Tucker

## Abstract

Correct floral development is a consequence of a sophisticated balance between environmental and molecular cues. Floral mutants provide insight into the main genetic determinants that integrate these cues, as well as providing opportunities to assess functional conservation across species. In this study, we characterize the barley (*Hordeum vulgare*) multiovary mutants *mov2.g* and *mov1* and propose causative gene sequences: a C2H2 zinc-finger *HvSL1* and a B-class gene *HvMADS16*, respectively. In the absence of *HvSL1,* flowers lack stamens but exhibit functional supernumerary carpels resulting in multiple seeds per floret when artificially pollinated. Deletion of *HvMADS16* in *mov1* causes homeotic conversion of lodicules and stamens into bract-like organs and carpels that contain non-functional ovules. Based on developmental, genetic, and molecular data we propose a model by which stamen specification in barley is defined by HvSL1 acting upstream of barley B-class genes, specifically the transcriptional up-regulation of *HvMADS16*. The present work identifies strong conservation of stamen formation pathways with rice, but also reveals intriguing species-specific differences. The findings lay the foundation for a better understanding of floral architecture in *Triticeae*, a key target for crop improvement.

**Highlight:** Analysis of the barley multiovary *mov1* and *mov2* loci indicates that HvSL1 and HvMADS16 exhibit both unique and conserved roles in the specification and development of cereal flowers.

## Introduction

The most diverse floral structures build upon similar molecular mechanisms, first described in the ABC model of flower development. This model, based on early studies in the dicotyledonous species *Arabidopsis thaliana*, *Antirrhinum majus* and *Petunia hybrida*, postulates that each organ within a flower is specified by the combinatorial action of specific gene classes. These are referred to as the A-, B-, C-, D- and E- class genes, which typically act in two adjacent whorls of the flower (Schwarz-Sommer *et al*., 1990; Coen and Meyerowitz, 1991; Colombo *et al*., 1995; Pelaz *et al*., 2000; Favaro *et al*., 2003). Generally, in the outermost whorl (whorl 1), A- and E-class genes control the development of sepals or other bract-like organs, while the action of A-, B- and E-class genes in whorl 2 gives rise to petals or equivalent structures. Stamens in the third whorl are specified by the function of B-, C- and E-class genes. In the fourth inner whorl, carpels require C- and E-class genes, while D-class genes are essential for ovule development. Most genes within the ABC model encode transcription factors belonging to the MADS-box gene family (Coen and Meyerowitz, 1991). In the thirty years since postulation of the ABC model, numerous studies have added to our knowledge, creating a more comprehensive view of the complex gene regulatory networks leading to flower formation (Thomson and Wellmer, 2018). Importantly, dimers of MADS-box proteins can interact together to form tetrameric complexes, described as the floral quartet model (Theissen, 2001). The floral tetramers can promote DNA looping to bring distal promoter regions closer and allow recruitment of transcription co-factors, chromatin remodelling proteins and other transcription factors to regulate the expression of downstream genes (Melzer and Theissen, 2009; Smaczniak *et al*., 2012).

Cys_2_/His_2_ (C2H2) zinc-finger transcription factors also play an important role in flower development by acting as upstream and downstream regulators of MADS-box genes, and by coordinating cell proliferation and differentiation during floral organogenesis (Lyu and Cao, 2018). In *Arabidopsis*, *SUPERMAN* (*SUP*) is involved in maintaining the floral organ boundary between whorls 3 and 4, as well as controlling the stamen, carpel and ovule primordia (Gaiser *et al*., 1995; Sakai *et al*., 1995; Breuilbroyer *et al*., 2016). *RABBIT EARS* (*RBE*) maintains the boundary between whorls 2 and 3, inhibits expression of the C-class gene *AGAMOUS* (*AG*) in the second whorl and causes formation of petal primordia (Takeda *et al*., 2004; Krizek *et al*., 2006). *KNUCKLES* (*KNU*) has been shown to regulate floral meristem determinacy and act downstream of *AGAMOUS* (Sun *et al*., 2009). Moreover, *JAGGED* (*JAG*) functions in forming the petal primordia and redundantly acts with its paralog *NUBBIN* (*NUB*) to control lateral growth and differentiation of stamens and carpels (Dinneny *et al*., 2004; Dinneny *et al*., 2006). In rice (*Oryza sativa*), the *JAG* orthologue *STAMENLESS1* (*SL1*) specifies lodicule and stamen identity (Xiao *et al*., 2009).

Understanding floral development provides information on how to manipulate floral structures for potential application in agriculture. In cereals, considerable effort has been devoted towards understanding flower development in rice, particularly in the context of the MADS-box genes (Arora *et al*., 2007; Yoshida and Nagato, 2011). Less information is available regarding members of the *Triticeae*, such as wheat (*Triticum aestivum*) and barley (*Hordeum vulgare*), despite their importance in the food and feed industries. Recent studies have mined genomic resources to address the diversity of ABC genes in barley (Callens *et al*., 2018; Kuijer *et al*., 2021). Coupled with the results of functional studies (e.g. *HvMADS1*, *HvLFY*, *HvAP2* and *HvMADS29*; Li *et al*., 2021; Selva *et al*., 2021; Shoesmith *et al*., 2021), a trend of functional diversification between temperate and tropical grasses is emerging. This may have implications for *Triticeae*-derived cereals in general, whereby manipulating floral structures to improve hybrid seed production and ultimately yield still represents a viable target for crop improvement (Selva *et al*., 2020). It also provides impetus to identify and characterise mutants in the *Triticeae* rather than relying solely on information derived from rice.

Diversity in floral structure might be introduced through specific targeting of known regulatory genes (e.g. CRISPR/Cas9; Selva *et al*., 2021; Li *et al*., 2021), through the exploitation of natural genetic variation in germplasm banks (e.g. in barley; Caldwell *et al*., 2004; Talamè *et al*., 2008; Druka *et al*., 2011; Szarejko *et al*., 2017; Szurman-Zubrzycka *et al*., 2018; Schreiber *et al*., 2019; and wheat; Krasileva *et al*., 2017), or through induced mutations in forward genetic screens. In terms of the latter, historical screens have uncovered a considerable array of phenotypes, some of which dramatically modify reproductive organ structure and function. One of these phenotypes is referred to as pistillody, the conversion of stamens into additional pistils (Murai and Tsunewaki, 1993; Murai *et al*., 2002; Peng 2003). Pistillody is one avenue to increase the number of seed-bearing units per plant (Yang and Tucker *et al*., 2021), as well as to create a male-sterile mother plant for cross-pollination in hybrid breeding (Selva *et al*., 2020). Studies in wheat mutants often show that pistillody correlates with expression changes of genes belonging to the ABC model (Yamada *et al*., 2009; Wang *et al*., 2015; Yang *et al*., 2015). This is particularly true for B-, C- and D-classes, which are directly implicated in stamen, carpel and ovule development.

Until now, at least three multiovary loci (*mov1*, *mov2* and *mov5*) have been reported in barley (Barley Genetics Newsletter, 2013). Mutations at these loci specifically affect flower development and cause additional carpels to grow at the total or partial expense of stamens. The barley *multiovary5* (*mov5*) locus was recently shown to encode a *FLORICAULA*/*LEAFY* (*HvLFY*) transcription factor on chromosome 2H (Selva *et al*., 2021). Phenotypic and molecular analysis of *HvLFY* revealed functional divergence relative to its orthologue from rice (*APO2*), particularly in relation to putative gene target sequences and a lack of impact on tillering (Selva et al., 2021). The *multiovary2* (*mov2*) locus, derived from a fast neutron irradiated population generated in the early 1990s (Soule *et al*., 1996), was roughly mapped to the short arm of chromosome 3H and suggested to encode a MADS-box gene (Soule *et al*., 2000). The *multiovary1* (*mov1*) locus was initially recorded as a homeotic conversion of stamens into additional non-functional carpels, with substitution of lodicules as half-curled leaf-like structures and partial fertility (Tazhin, 1980). Linkage and mapping attempts of *mov1* alleles in the barley varieties Revelatum (spontaneous mutation) and Steptoe (fast- neutron mutagenesis) positioned the locus on the centromeric region of chromosome 7H and it was proposed this locus may also encode a member of the MADS-box gene family (Tazhin, 1980; Soule *et al*., 1996; Soule *et al*., 2000).

Despite an early general description of the mutant phenotype and rough mapping of the *mov2* and *mov1* loci, the causative gene sequences have yet to be identified. Apart from rice, little is known about flower development in domesticated grass species. To widen our knowledge of floral development in the temperate grasses, this study aimed to map and characterise the barley *mov1* and *mov2* loci. We show that *mov2* maps to a region on 3H encoding a C2H2 zinc-finger transcription factor, named here as *HvSL1* based on homology to rice *STAMENLESS1* (Xiao *et al*., 2009). Moreover, we show that a total of three genes on chromosome 7H, including the MADS-box B-class gene *HvMADS16* are absent in *mov1* plants. We investigate the interaction of HvSL1 with barley B-class genes and report the developmental changes and effect on ABC gene expression in *mov1* and *mov2.g* during floral development. We explore the implications of *HvMADS16* absence in the context of flower development and postulate a model for stamen specification in barley. Our results highlight intriguing differences between cereals in terms of MADS-box dependent molecular regulation, as well as strong conservation with rice in the core determinants of stamen formation. This study also establishes *Triticeae*-specific resources that may prove useful in future attempts to modify floral architecture for breeding.

## Materials and Methods

### Plant material and genotyping

Seeds segregating for the *mov2.g* allele (*mov2* locus) and for *mo6b* allele (*mov1* locus) in cv. Steptoe were kindly provided by Prof. A. Kleinhofs. Germination was synchronised by cold treatment at 4°C for three days in the dark. The germinated seeds were then transplanted into pots and grown in the glasshouse at 23°C/16°C day/night temperatures and long days (∼12 hours). Phenotyping was done at heading stage by visually inspecting the central florets of several spikes per plant. Florets were imaged with a Nikon SMZ25 Stereo Fluorescence Microscope equipped with DS-Ri1 colour cooled digital camera.

Genotyping of the plants was performed by copy number analysis using TaqMan^TM^ assay (Thermo Fisher Scientific, USA) designed for *HvSL1* and *HvMADS16* as genes of interest. The barley *CONSTANS*-like *CO2* (*HORVU6Hr1G072620*) was used internal positive control (Kang *et al*., 1997). The sequence of primers and probes can be found in Supplementary **Table S1A** at *JXB* online. The reaction was set up as follows: 1x of PrecisionFast^TM^ qRT- PCR Master Mix with Low ROX (Primerdesign Ltd., UK), 200 nM of each primer, 100 nM probe, 150 ng template DNA and water to a final volume of 10 μL. The reaction was performed with an initial activation step at 95 °C for 2 minutes, followed by 40 cycles at: 95 °C for 5 seconds and 60 °C for 20 seconds in a QuantStudio 6 Flex Real-Time PCR machine (Thermo Fisher Scientific, USA). The detectors used were FAM-BHQ1 and HEX- BHQ1, with ROX as internal passive reference.

The *mov2* bi-parental mapping population was created by artificially pollinating *mov2.g* flowers (cv. Steptoe) with wild-type pollen from cv. Morex. Heterozygosity of F_1_ plants was confirmed by KASP^TM^ marker analysis across two known SNPs, at positions chr3H_1006543 and chr3H_28805649 and by Sanger sequencing across a third SNP at position chr7H_557244517 according to the Morex reference assembly (Hv_IBSC_PGSB_v2). For fine-mapping of the *mov2* locus, SNPs spanning the 3HS region were identified and developed as KASP^TM^ markers. SNPs were identified from a published Steptoe x Morex dataset (Close *et al*., 2009) and from Steptoe leaf transcriptomic data mapped to the reference Hv_IBSC_PGSB_v2 Morex assembly. Primer Picker (LGC Genomics) and the LGC Genomics SNPline^TM^ were used to design and prepare the KASP™ marker assays. Assays were performed using KASP^TM^ Master mix as instructed by the manufacturer. Sequence of KASP^TM^ markers can be found in **Table S1B**.

### Nucleic acid extraction and genomic PCR

For genotyping, genomic DNA was extracted from freeze-dried 2-weeks old plant material using a phenol/chloroform method as described by Rogowsky *et al*., 1991. PCR was performed using Q5® high-fidelity DNA polymerase (New England BioLabs, USA), following the manufacturer’s protocol in a final volume of 12.5 μL. All PCR products were separated by agarose gel electrophoresis, purified using ISOLATE II PCR and Gel Kit (Bioline, Australia) and Sanger sequenced at the Australian Genome Resource Facility (AGRF). Primers used for PCR can be found in **Table S1C, 1D, 1E**.

### RNA extraction, cDNA synthesis and quantitative real-time PCR (qRT-PCR)

Ultra fine-pointed tweezers were used to manually dissect inflorescences at developmental stages W2.0, W3.5, W4.0, W6.0 corresponding approximately to 17, 20, 23 and 26 Days Post Germination (DPG) in the growing conditions described above. Inflorescences were snap-frozen in liquid nitrogen, RNA extraction was performed using the ISOLATE II RNA Plant Kit (Bioline, Australia), followed by TURBO^TM^ DNase treatment (Thermo Fisher Scientific, USA) and cDNA synthesis using SuperScript^TM^ IV First Strand Synthesis (Thermo Fisher Scientific, USA) as per manufacturers’ instructions.

RNA from transfected protoplasts was extracted using a phenol-chloroform method. Briefly, samples were lysed in a 1.5 mL tube and homogenized in 500 μL TRIzol (Thermo Fisher Scientific, USA) before adding 150 μL chloroform. Samples were vortexed and centrifuged at maximum speed at 4°C for 15 minutes. The aqueous phase was carefully transferred to a new 1.5 mL tube with 250 μL isopropanol and incubated at 4°C for 10 minutes before centrifuging at maximum speed at 4 °C for 20 minutes. The precipitated RNA was washed with 500 μL 100% ethanol, allowed to air-dry and resuspended in 20 μL DEPC-treated water.

Quantitative real-time PCR (qRT-PCR) was carried out as described in Selva *et al*., 2021. Each timepoint is the result of three technical replicates and at least three biological replicates. Absolute quantification is reported as normalised transcript against the geometric means of the three least varying control genes as described in Vandesompele *et al*., 2002. Relative quantification in transfected protoplasts is expressed as calibrated Normalised Relative Quantities (NRQ) as calculated by the qbase+ software (Biogazelle, Belgium – Hellemans *et al*., 2007). Primer information for the reference genes and genes of interest can be found in **Table S1F**.

### CRISPR and plant transformation

Guide RNA design and cloning for *HvSL1* CRISPR knockout was done following the procedure described by Ma *et al*., 2015 with vectors kindly provided by Prof Yao-Guang Liu (South China Agricultural University). Two guide RNAs (gRNA) targeting positions +42 bp and +275 bp from the translational start site of *HvSL1* were cloned in the same vector. Primer sequences used for cloning are listed in **Table S1G**.

*Agrobacterium tumefaciens*-mediated plant transformation (cv. Golden Promise) and genotyping of regenerant plants was performed as described in Selva *et al*., 2021.

### In situ hybridization

Sense and antisense RNA probes for *in situ* hybridization were amplified using Q5® high-fidelity DNA polymerase (New England BioLabs, USA) with the T7 promoter extension to the 5’ of primers (primers used can be found in **Table S1H**), transcribed and DIG-labelled. Inflorescences were prepared and *in situ* hybridization was performed as described in Selva *et al*., 2021.

### Bi-molecular fluorescent complementation (BiFC)

The full-length coding sequences of *HvMADS2*, *HvMADS4*, *HvMADS16* and Δ*HvMADS16,* containing only the MADS-domain (aa1-65), were PCR-amplified with Q5® high-fidelity DNA polymerase (New England BioLabs, USA) and cloned in the BiFC vectors pSAT1-nEYFP-N1 (N-terminal fragment) and pSAT1-cEYFP-C1-B (C-terminal fragment) (Citovsky *et al*., 2006). Primers used for cloning are listed in **Table S1I**. All plasmids were checked by digestion and Sanger sequencing. Biolistic particle bombardment of onion epidermal cells was carried out as described in Selva *et al*., 2021.

### Light and electron microscopy

For light microscopy, wild-type, *mov2* and *mov1* carpels were fixed in FAA solution (50% ethanol 100%, 5% acetic acid, 10% formaldehyde 37%, one drop of Tween-20) overnight, dehydrated in a 70-100% ethanol series and embedded in LR white resin. Samples were sectioned using a Leica Rotary Microtome RM2265 at 1.5 μm. Slides were stained with 0.1% toluidine blue in 0.1% sodium tetraborate for 2 minutes and rinsed three times with water, dried and mounted with DPX. After 72 hours slides were imaged with a Nikon Eclipse Ni-E optical microscope equipped with DS-Ri1 colour cooled digital camera. Image analysis and processing was carried out with the NIS-Elements AR software.

For scanning electron microscopy (SEM), inflorescences were manually dissected and fixed overnight in 4% paraformaldehyde, 1.25% glutaraldehyde in PBS, 4% sucrose, pH 7.2. Before processing, samples were washed three times in PBS and fixed in 2% OsO_4_ in PBS for one hour. Samples were then dehydrated in a 50-100% ethanol series and dried with a critical point dryer. Dried samples were arranged on carbon tabs stuck to 12 mm aluminium stubs and coated with platinum. Samples were observed using a Hitachi FlexSem 1000 Scanning Electron Microscope.

### Protoplast isolation

Isolation of barley leaf protoplasts was performed as described by Yoo *et al*., 2007 with minor modifications. Briefly, the adaxial epidermal layer of leaves from 11-day old barley seedlings (cv. Golden Promise) was manually peeled off and the leaf was cut into approximately 2 cm x 0.5 cm strips using surgical blades. The leaf segments of approximately 10 plants were immediately transferred in a Petri dish containing 15 mL of 0.6 M mannitol for 30 minutes at room temperature to induce plasmolysis. After incubation, the leaf segments were transferred to another Petri dish containing 10 mL of freshly prepared enzyme solution [0.55 M mannitol, 40 mM MES-KOH at pH 5.7, 20 mM KCl, 2.0% cellulase R10 (Yakult Pharmaceutical, Japan), 0.75% macerozyme R10 (Yakult Pharmaceutical, Japan), 10 mM CaCl_2_, 0.1% BSA] and incubated for 3 hours in the dark at 28 °C with gentle shaking (40-60 rpm). After enzymatic digestion, forceps were used to gently remove the remaining epidermis and leaf debris from the enzyme solution. An equal volume (10 mL) of W5 solution (154 mM NaCl, 125 mM CaCl_2_, 5 mM KCl and 2 mM MES-KOH at pH 5.7) was slowly added to the protoplasts and the solution was filtered with a 100 μM nylon mesh to a 50 mL round-bottom tube. Volume was adjusted by adding 5 mL of W5 solution. The filtered protoplasts were collected by centrifugation at 600 g for 3 minutes. Supernatant was replaced with 15 mL of fresh W5 and the protoplasts resuspended by gentle shaking. Protoplasts were allowed to pellet by gravity for 30 minutes in ice. After incubation, the supernatant was promptly removed and substituted with MMG solution (0.6 M mannitol, 15 mM MgCl and 4 mM MES-KOH at pH 5.7) at a concentration of 10^6^ cells/mL, determined by counting cells in 12 μL of a 1:10 dilution of protoplast solution with a haematocytometer.

### PEG-mediated transfection of barley protoplasts

PEG-mediated transfections were mostly carried out as described by Bai *et al*., 2014. Firstly, 200 μL of PEG-Ca^2+^ solution [40% (w/v) PEG 4000, 0.4 M mannitol, 0.1 M CaCl_2_] were pre- loaded in pipette tips. Secondly, 100 μL of protoplast solution (approximately 10^5^ cells) were added to 20 μg of each plasmid DNA in a 2.0 mL tube. The pre-loaded PEG-Ca^2+^ solution was immediately added to the protoplast-DNA mixture, mixed gently and incubated for 15 minutes at room temperature in the dark. The transfection process was stopped by adding 840 μL of W5 and centrifuged at 600g for two minutes. Cells were resuspended in 500 μL W5 and transferred to multi-well plates previously coated with 5% (v/v) FBS. Protoplasts were cultured at 28 °C for 40 - 48 hours in the dark.

To test transfection efficiency, protoplasts were transfected with pUbi-YFP-rbcS in six independent transfections. 40-48 hours after transfection the protoplasts were visualized under UV and bright light using a Nikon Eclipse Ni-E optical microscope equipped with DS- Ri1 colour cooled digital camera.

## Results

### mov2.g florets have functional supernumerary carpels and produce multiple seeds

In wild-type barley, floral organs are arranged in a defined pattern such that the outermost whorl contains the palea and lemma (whorl 1). These are followed by two lodicules in whorl 2 and three stamens in whorl 3, surrounding a single carpel at the centre of the floret in whorl 4 (**Fig. 1A, B**) (Brenchley, 1920).

**Fig. 1.**
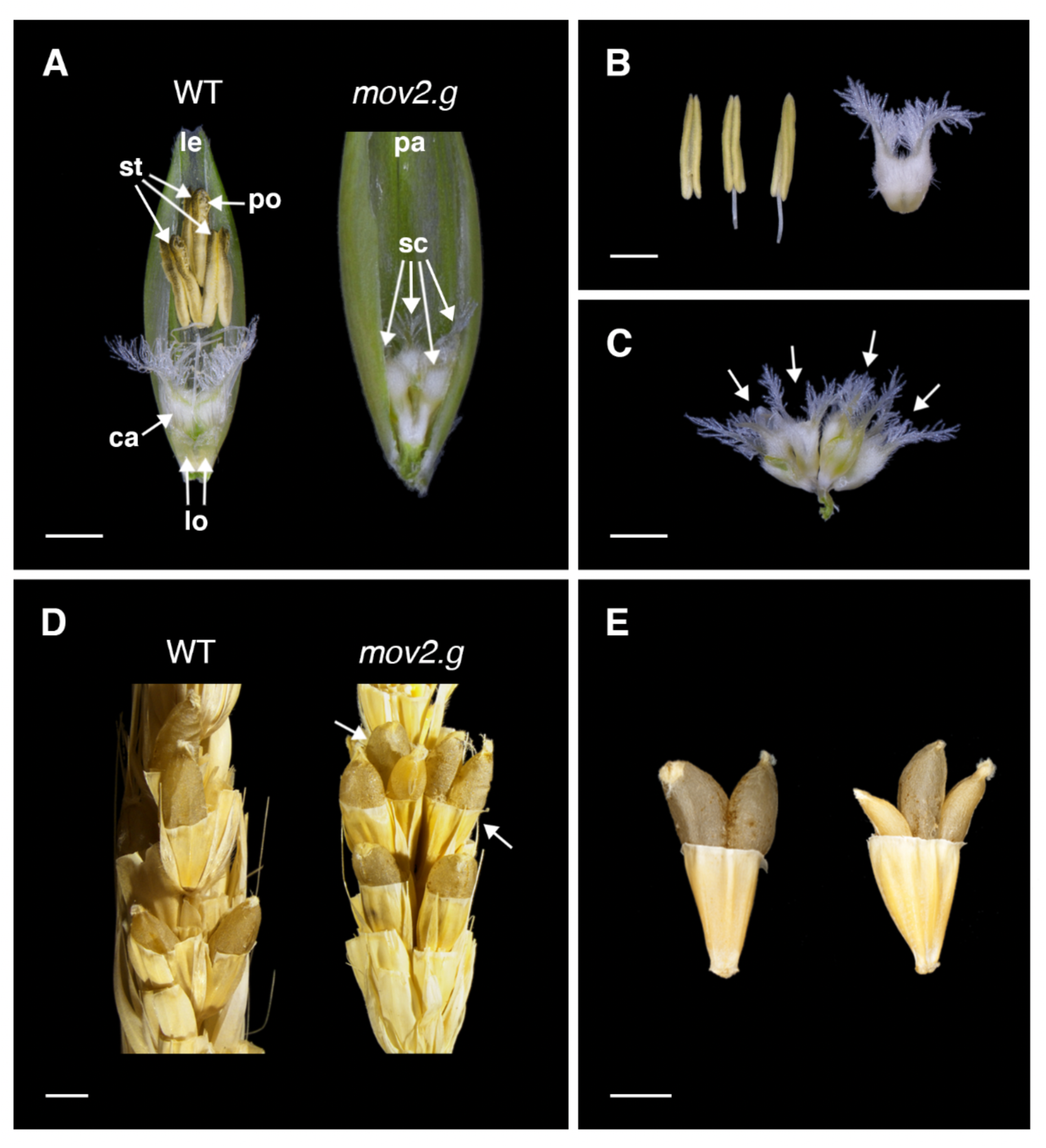
Florets, reproductive organs and seeds in wild-type and *mov2.g*. (**A**) Exposed wild-type (WT) and *mov2.g* florets. Palea or lemma have been removed to show internal floral organs. (**B**) Reproductive organs in wild-type flowers consist of 3 stamens and 1 carpel. (**C**) Reproductive organs in *mov2.g* flowers consist of supernumerary carpel-like structures (arrows). Scale bars: 1000 µm. (**D**) Artificially pollinated wild-type (WT) and *mov2.g* spikes. White arrows indicate multiple seeds per floret. (**E**) Examples of multiple seeds per floret produced by artificially pollinated *mov2.g* spikes. Scale bars: 2000 µm. Lemma (le), palea (pa), lodicule (lo), stamen (st), carpel (ca), supernumerary carpel-like structure (sc), pollen (po).

In florets from *mov2.g* plants, the stamens are replaced by supernumerary carpels rendering the plant unable to self-pollinate (**Fig. 1A**). The number of carpels within each floret typically varies between 5 - 7 carpels or carpel-like structures. Visually, all carpels appear irregularly shaped, joined at the base, and of smaller size relative to a wild-type carpel (**Fig. 1C**). In contrast, development and morphology of the lemma, palea and lodicules remains largely unaffected. Most notably, when *mov2.g* plants were used as a female recipient in artificial pollination, most florets were able to produce multiple seeds with a maximum of three developing seeds per floret (**Fig. D, E**). This is due to the ability of some *mov2.g* carpel structures to correctly differentiate fully functional embryo sacs, as was observed in transverse sections stained with toluidine blue (**Fig. S1**). All seeds from *mov2.g* plants were viable, and after germination gave rise to mature plants.

To better understand the developmental basis of the multiovary phenotype, scanning electron microscopy (SEM) was used to compare wild-type (cv. Steptoe) and *mov2.g* inflorescences. Immature inflorescences were morphologically similar at the very early stages of development, corresponding to floret primordium (Waddington stage W3.0) (**Fig. 2**) (Waddington and Cartwright, 1983). At stages W3.5 – 3.75, both wild-type and *mov2.g* floral meristems initiated lateral dome-shaped protrusions, consistent with the appearance of stamen primordia from the floral meristem. Following this stage, the first morphological differences were identified. In wild-type, the stamen primordia differentiated into filament and anther tissues (W5.0 – 8.5), while the meristematic tissue at the centre proliferated and terminally differentiated into a single ovule-containing carpel. As each wild-type floral meristem developed, they maintained a vertical symmetry along the central inflorescence rachis.

**Fig. 2.**
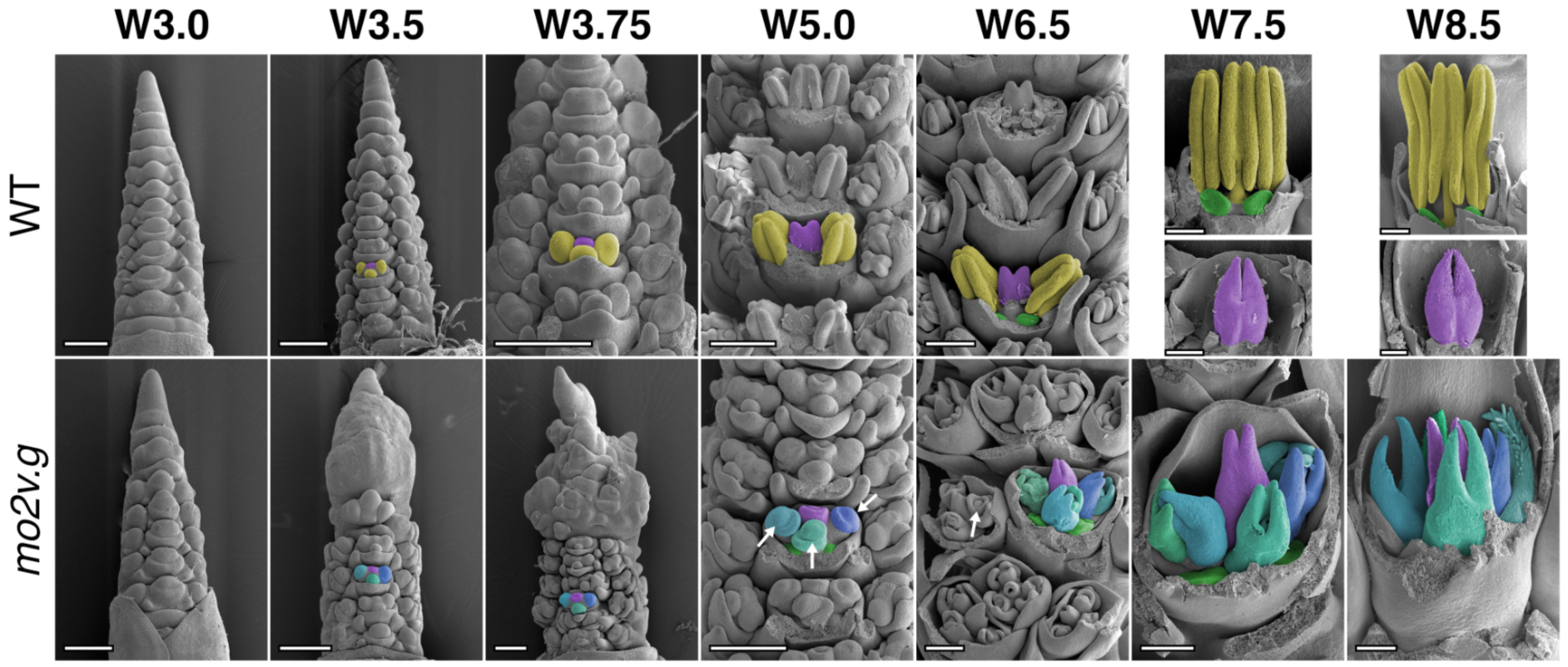
Scanning electron microscopy (SEM) of *mov2.g* developing inflorescences. In wild-type (WT), primordia giving rise to stamens are false-coloured in yellow, cells giving rise to the carpel in purple and lodicules in green. In *mov2.g*, the central carpel (purple) and lodicules (green) appear to be retained, cells giving rise to the additional carpel-like structures are false-coloured in blue (different shades). White arrows indicate ovule primordia. Waddington stage is indicated for each developmental timepoint, from stage W5.0 lemma and/or stamens have been removed to expose the carpels. Scale bars: 200 µm.

In contrast, in *mov2.g* the lateral protrusions did not differentiate into stamens, but instead gave rise to organs that followed the characteristic morphogenesis of carpels (W5.0 – 8.5) (**Fig. 2**). Specifically, the carpel-like organs appeared to initially develop an ovule primordium that was later surrounded by the growing carpel tissue. By stage W8.5, the additional carpel-like structures also differentiated stigmas bearing stigmatic papillae and appeared to develop synchronously with the central carpel, resulting in a unit of multiple carpel-like structures joined basally. Moreover, as floral organs developed there was frequent loss in the vertical symmetry of florets along the central inflorescence rachis and fasciation of the growing tip, which was then reflected in a shorter and broader spike compared to wild-type (**Fig. S2**).

### mov2 encodes a C2H2 zinc-finger transcription factor

A previous study had mapped the barley *mov2* locus to the telomeric region on the short arm of chromosome 3H (3HS) within a ∼ 28 Mb genomic interval (Soule *et al*., 2000). This interval is too large to reliably identify the underlying causative gene(s). Therefore, a *mov2*.g (cv. Steptoe) x Morex bi-parental population was developed to map *mov2* to a higher genetic resolution. The multiovary phenotype was not observed in F_1_ individuals, indicative of the *mov2*.g mutation being completely recessive. A total of 352 F_2_ plants were grown until maturity and used for recombination-based genetic mapping.

Seven KASP^TM^ markers designed to span the previously published interval confirmed that *mov2* mapped to the expected region on chromosome 3H. Following this, an additional 10 KASP^TM^ markers were designed to saturate the interval, reducing the critical *mov2*- containing region to approximately 1.9 Mb, based on flanking markers chr3H_9095799 and chr3H_11039299 (**Fig. S3**). Further resolution of the locus was achieved with six additional KASP^TM^ markers in 179 F_3_ individuals segregating for the multiovary phenotype. Comparison of phenotypic and genotypic segregation across these F_3_ plants reduced the critical interval to roughly 449 Kb, between markers chr3H_9748112 and chr3H_10289104 (**Fig. S3**). This region contains 20 annotated gene sequences, based on the Morex Scaffold_1432 (*personal communication* Dr. Martin Mascher, IPK Gatersleben, Germany) (**Table S2**).

Expression in floral tissues was assessed for each of the 20 annotated gene sequences to identify the likely causative gene, especially in tissue types affected in *mov2.g* florets. Barley expression datasets were obtained from a range of transcriptomes including Steptoe and *mov2.g* leaves (Zadoks stage Z22; Zadoks *et al*., 1974), 2-week old seedlings (Z12; Liu *et al*., 2019), developing inflorescences at stages W2.0, W3.5 and W8.0 - 8.5 (Liu *et al*., 2019), and developing pistils (W8.0 – 10.0; Matros *et al*., 2021). Overall, ten of the 20 annotated gene sequences showed expression in either pistils or inflorescences. Of these, only *HORVU3Hr1G003740* was identified to be uniquely expressed in reproductive tissues but not in vegetative tissues (leaf and seedling stage) nor in *mov2.g* leaves (**Fig. S4**). Sequence analysis indicated that *HORVU3Hr1G003740* encodes a putative C2H2 zinc-finger transcription factor sharing 65.6% sequence identity with the rice protein STAMENLESS1 (OsSL1) (**Fig. S5**). In rice, *OsSL1* is known to play a crucial role in floral development, with loss of *OsSL1* function leading to a multiovary phenotype (Xiao *et al*., 2009). Thus, *HORVU3Hr1G003740* is hereafter referred to as *HvSL1*.

To validate the transcriptomic data, mRNA *in situ* hybridization was performed to confirm *HvSL1* spatial expression pattern in wild-type developing inflorescences. At stage W3.0, *HvSL1* expression accumulated in the floret primordia, including the inflorescence tip, and was progressively found in the primordia of glumes, lemma, lodicules, stamens, carpels and ovules (W3.5-5.0) (**Fig. 3**). As the floral organs developed (W7.0) the signal persisted in the stamens, in the carpel, at the apical tip and central vasculature of the lemma but had decreased in the lodicules.

**Fig. 3.**
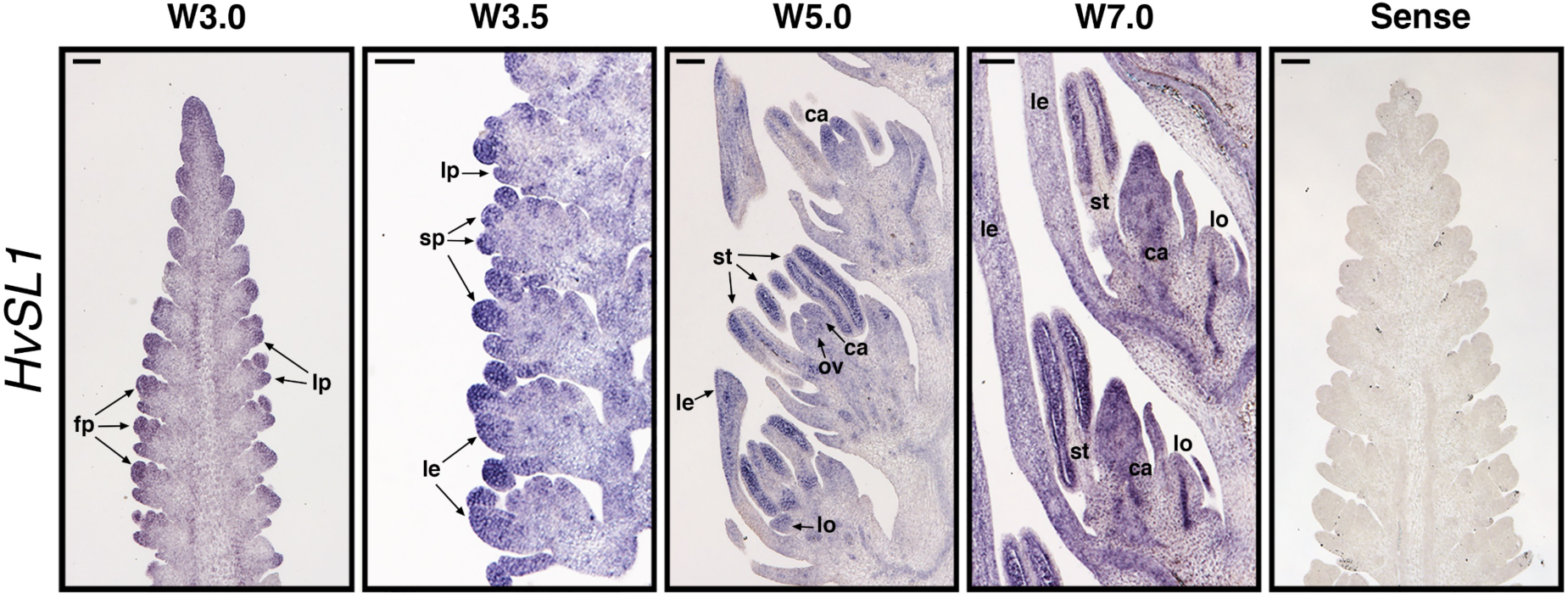
Spatial expression of *HvSL1* during wild-type inflorescence development. Waddington stages correspond to lemma-floret primordia (W3.0), stamen primordia (W3.5), ovule primordium (W4.5) and stamen and carpel development (W7.0). Annotations and arrows indicate floral primordia (fp), lemma primordia (lp), stamen primordia (sp), lemma (le), lodicule (lo), stamen (st), carpel (ca) and ovule (ov). A sense *HvSL1* probe was used as negative control to determine probe specificity. Scale bars: 100 µm.

### HvSL1 is deleted in mov2.g plants and its absence causes multiovary

To determine whether the *mov2.g* phenotype is a consequence of a loss-of-function in *HvSL1* and not of other gene sequences in the genetically mapped interval, PCR primers were designed to amplify the entire *HvSL1* coding sequence. Comparisons between Morex, Steptoe and *mov2.g* (cv. Steptoe) genomic DNA suggested that *HvSL1* is absent in *mov2.g* plants but present in both Steptoe and Morex (**Fig. S6A**). By contrast, predicted high-confidence neighbouring gene sequences were still present, indicating that *HvSL1* could be the only deleted gene at the *mov2* locus (**Table S2**). For additional confirmation, the *HvSL1*- specific PCR was repeated on 36 critical recombinant F_3_ individuals from the *mov2.g* x Morex mapping population. For all samples, the absence of *HvSL1* correlated with mutant phenotype expression. Although this alone does not provide conclusive proof, it appears highly likely that the multiovary phenotype in *mov2.g* is due to deletion of the *HvSL1* gene. Furthermore, *HvSL1* expression by quantitative real-time PCR (qRT-PCR) in developing inflorescences indicated that *HvSL1* expression was completely absent in *mov2.g* samples (**Fig. S6B**), whereas *HvSL1* transcript abundance steadily increased in wild-type between stages W2.0 - W4.5 (double ridge – carpel primordium) with a slight decrease only at stage W6.0 (carpel development).

To provide additional support for the hypothesis that *HvSL1* is the causative gene sequence for the *mov2.g* phenotype, *Hvsl1* knockout plants were generated by CRISPR-Cas9 editing. Transformation of immature barley embryos (cv. Golden Promise) led to the generation of five T_0_ lines – HvSL1-1, HvSL1-3, HvSL1-5, HvSL1-6 and HvSL1-10 – that phenocopied *mov2.g* plants. Specifically, HvSL1-6 was found to be a homo-allelic edit while HvSL1-1, HvSL1-3, HvSL1-5 were all bi-allelic and HvSL1-10 was hetero-allelic (**Fig. S7**). In all cases, edits occurred at the canonical cut site and resulted in loss-of-function frame-shift mutations. The resulting phenotypes differed among lines reflecting variation in phenotypic expression as was also observed in *mov2.g* plants. This variation was observed in the degree of homeotic conversion of stamens to carpels, particularly in the number and complexity of supernumerary carpel-like structures, while all retained correct lodicule development (**Fig. 4**).

**Fig. 4.**
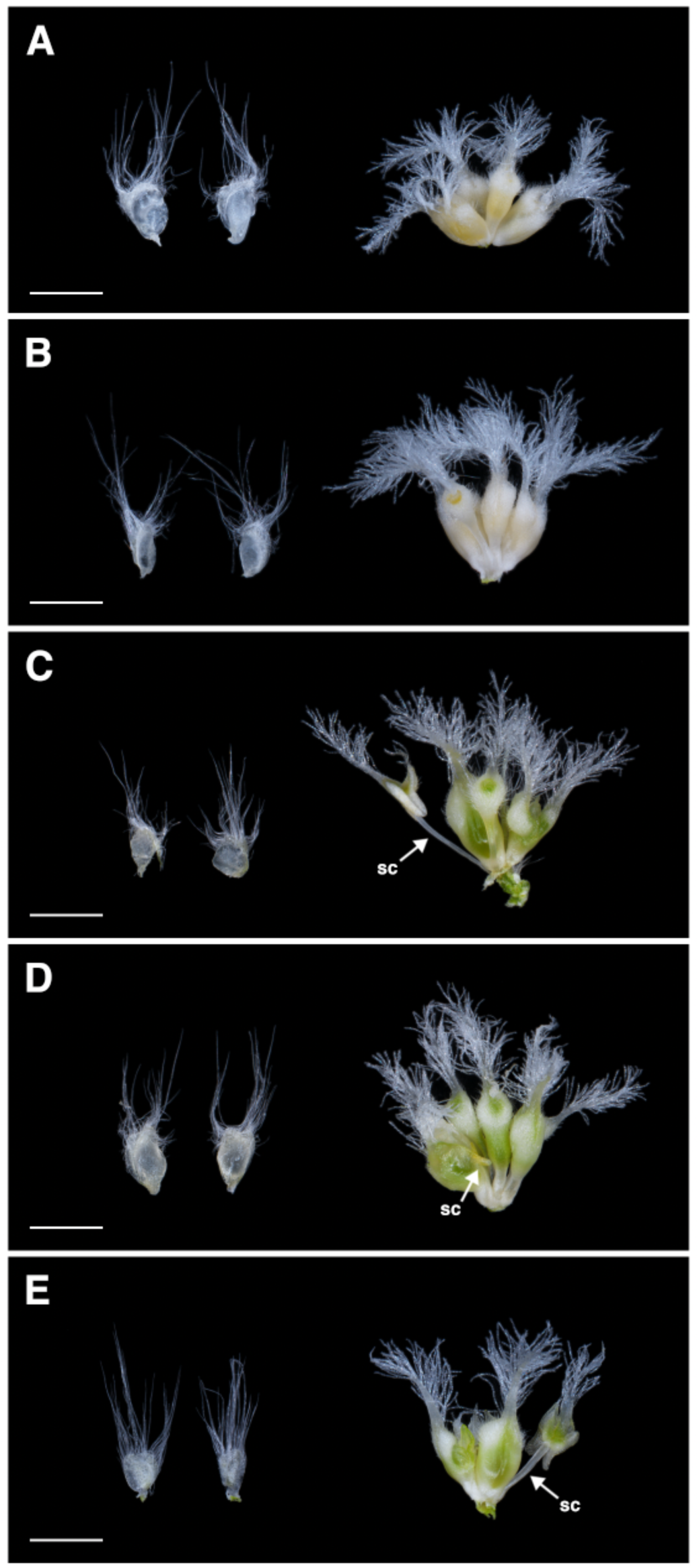
Phenotype of CRISPR *Hvsl1*-knockout plants. All knock-out plants contained 2 lodicules and supernumerary carpels. (**A**) HvSL1-1, (**B**) HvSL1-3, (**C**) HvSL1-5, (**D**) HvSL1-6 and (**E**) HvSL1-10. Partial stamen conversions (sc) are annotated. Scale bars: 100 µm.

### HvSL1 influences HvMADS16 expression but not that of other B-class genes

Since the early 1990s flower development has always been described in the framework of the ABC model, whereby specific classes of genes are needed for the correct specification of each floral organ (Coen and Meyerowitz, 1991). The abnormal floral development in *mov2.g* is restricted to stamens and carpels consistent with altered activity of B-class genes. We therefore aimed to evaluate barley MADS-box B-class genes known to specifically act in stamen development. Using a homology-based search, three B-class genes were found in barley, consisting of 2 GLO-like homologues (referred to here as *HvMADS2* and *HvMADS4*) and a single DEF-like homologue (referred to as *HvMADS16*), confirming already published data (Callens *et al*., 2018).

B-class proteins act in heterodimeric complexes involving DEF and GLO proteins (Winter *et al*., 2002). To assess interactions among the barley B-class proteins, bimolecular fluorescence complementation (BiFC) was conducted in onion epidermal cells. Fluorescence signal was observed only in cells co-expressing *HvMADS16* with *HvMADS2* or *HvMADS4* (**Fig. S8**). For these combinations, fluorescence was predominantly confined to the nucleus. No signal was observed when the two GLO-like genes *HvMADS2* and *HvMADS4* were co-expressed in the same cells, or with a truncated version of *HvMADS16* (Δ*HvMADS16- nYFP*), containing only the MADS-domain. Likewise, no signal could be detected when homodimeric combinations for each B-class gene were tested (**Fig. S8**).

To determine whether the heterologous protein interaction assays are consistent with gene expression patterns, the expression domain of the B-class genes was assessed by *in situ* hybridization in developing inflorescences and flowers (**Fig. 5**). In wild-type (cv. Steptoe), *HvMADS2* was found to be expressed in most floral tissues (glumes, lodicules, stamen primordia and carpel) in the early stages (W3.5-6.0), but the signal became more specific as development progressed (**Fig. 5A**). In stamens, expression was initially localised mainly in the locules, but was later restricted to the filament (W7.0), as seen also for *HvMADS4* (**Fig. 5B**). However, *HvMADS4* was also observed to accumulate in the ovule primordium (W4.5- W6.0). At maturity, both *HvMADS2* and *HvMADS4* were detected at the apex of the carpel, as well as in the lodicules. Expression of *HvMADS16* was first detected in the stamen and lodicule primordia (W3.5) (**Fig. 5C**). As the stamens develop, *HvMADS16* signal became weaker, while remained strong in the lodicules (W6.0-W7.0). No expression was observed in the carpel or ovule at any stage and no signal was detected with the sense probes.

**Fig. 5.**
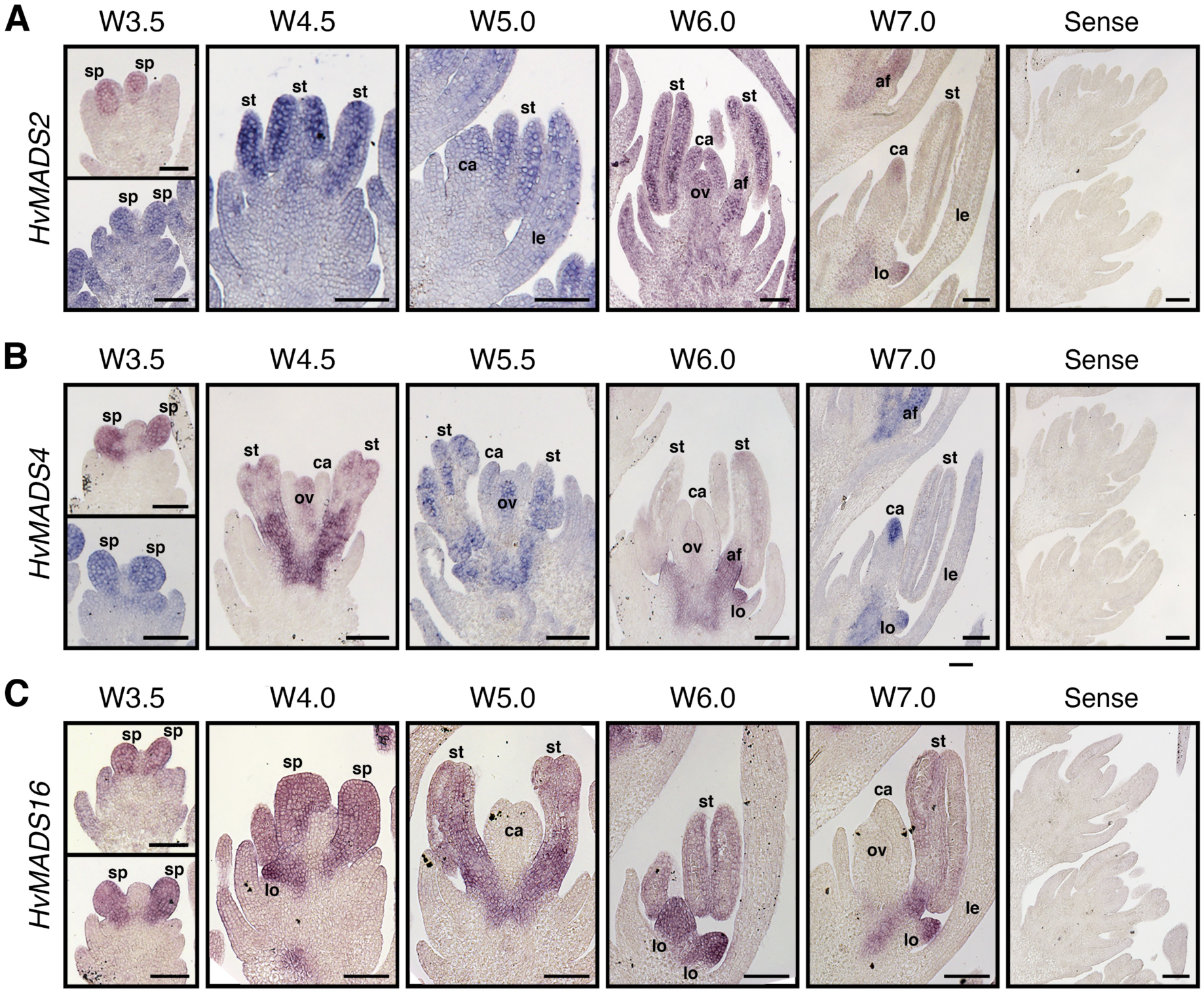
Expression pattern as assessed by *in situ* hybridization for barley B-class genes. (**A**) *HvMADS2*, (**B**) *HvMADS4* and (**C**) *HvMADS16*. Annotations indicate stamen primordia (sp), lemma (le), lodicule (lo), stamen (st), anther filament (af), carpel (ca) and ovule (ov). Scale bars: 100 µm.

The overlapping expression domain of *HvSL1* and B-class genes, together with the specific *mov2.g* phenotype suggest that these genes may act in a common pathway. Moreover, studies conducted in rice suggest that OsSL1 acts as an upstream positive regulator of *OsMADS16* transcription (Xiao *et al*., 2009). Expression assays in isolated barley protoplasts were performed to investigate the interaction between HvSL1 and barley B-class genes (**Fig. S9**). Independent transfection experiments showed that protoplasts carrying a *p35S:HvSL1* construct showed higher *HvSL1* expression relative to protoplasts transfected with a mock solution (**Fig. S9B**). In addition, a qRT-PCR assay indicated that endogenous *HvMADS16* transcript abundance was significantly increased in transfected protoplasts compared to the mock, while *HvMADS2* and *HvMADS4* expression remained unchanged (**Fig. S9C**). This suggests that HvSL1 may act as a positive regulator of *HvMADS16*.

### The B-class gene HvMADS16 is absent in mov1 plants

To further dissect the relationship between HvSL1 and *HvMADS16* we aimed to create a *Hvmads16* mutant using CRISPR/Cas9. Despite multiple attempts with different guides, edited plants were not recovered. To overcome this, we investigated historical induced mutant resources again. At least three multiovary loci have been reported to date (Barley Genetics Newsletter, 2013). Of these, the *multiovary1* (*mov1*) locus (Tazhin, 1980; Soule *et al*., 2000) bears a striking resemblance to the rice *Osmads16/superwoman1* mutant (Xiao *et al*., 2009). In barley *mov1* florets, lodicules are transformed into one pair of leaf-like organs and stamens show homeotic conversion into additional carpels (**Fig. 6**). In contrast, the central carpel in the fourth whorl is retained, as well as the palea and lemma. Unlike *mov2.g/Hvsl1* mutants, even if *mov1* florets develop a total of four carpels and each carpel lobe contains an ovule-like structure, these organs are sterile as the female gametophyte fails to differentiate correctly (**Fig. S10**). Furthermore, *mov1* florets did not produce any seeds when artificially pollinated with wild-type pollen.

**Fig. 6.**
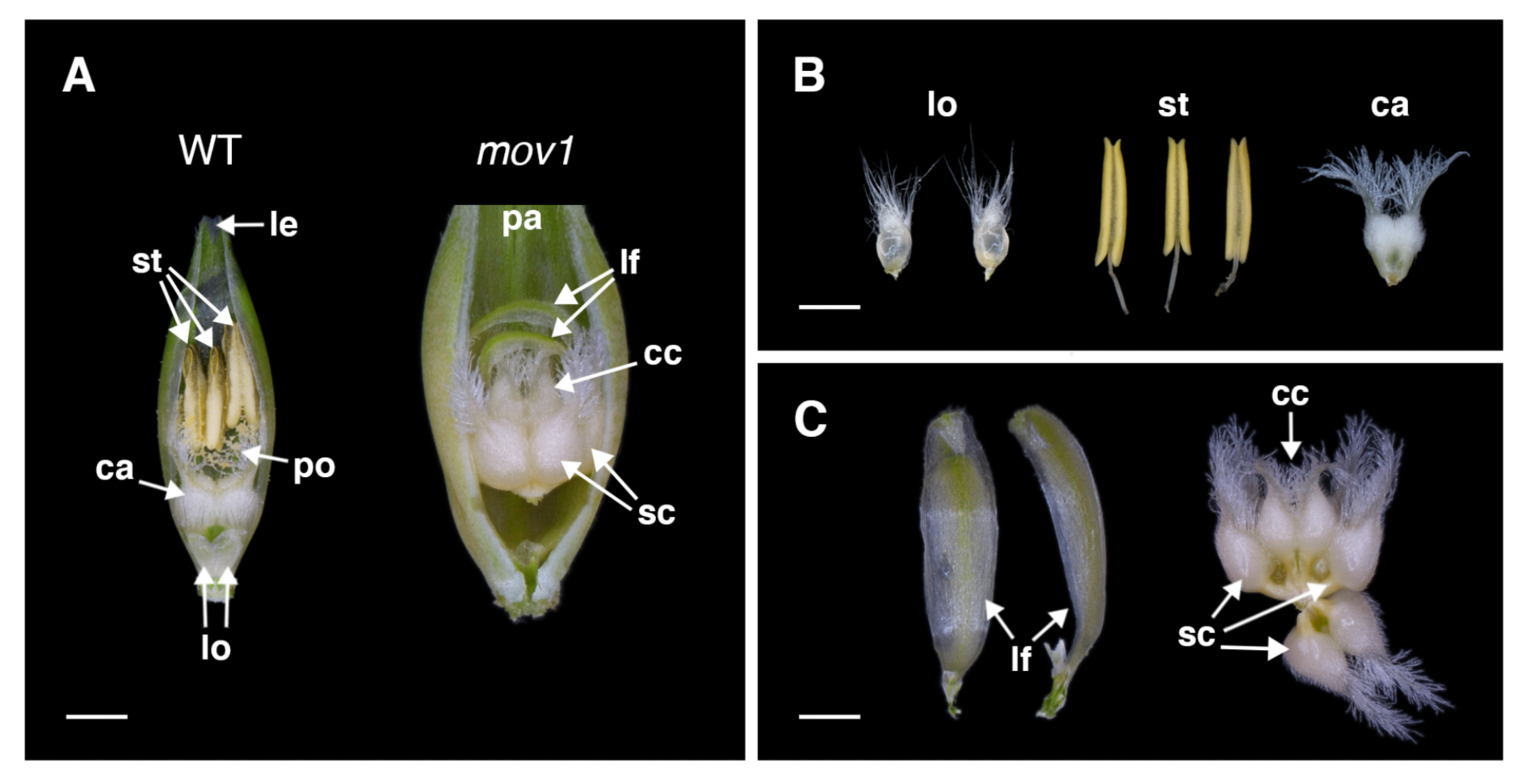
Florets and floral organs in wild-type and *mov1*. (**A**) Exposed wild-type (WT) and *mov1* florets. Palea or lemma have been removed to show internal floral organs. (**B**) The floral organs in a wild-type flower consist of 2 lodicules, 3 stamens and 1 carpel. (**C**) In *mov1* flowers, the lodicules are converted into bract-like organs, and stamens are converted into additional carpels. Scale bars: 1000 µm. Lemma (le), palea (pa), lodicule (lo), stamen (st), carpel (ca), supernumerary carpel-like structure (sc), central carpel (cc), leaf-like organ (lf), pollen (po).

SEM images of wild-type and *mov1* developing inflorescences were compared to determine the effect of *mov1* on floral organ development (**Fig. 7**). The earliest observable difference was seen immediately preceding the appearance of stamen primordia at Waddington stage W3.0, whereby a crease appears in the basal floral meristems of *mov1* inflorescences. At stage W3.5-3.75, instead of developing dome-shaped stamen primordia, the meristems in *mov1* divide into irregularly shaped protrusions which arrange into discernible multiple concentric creases by stage W5.0. As development progresses (W6.5-7.0) each crease develops into a carpel eventually leading to the 4-carpel structure visible in the mature *mov1* floret. Occasionally, it was observed that a single floral meristem could give rise to two distinct florets.

**Fig. 7.**
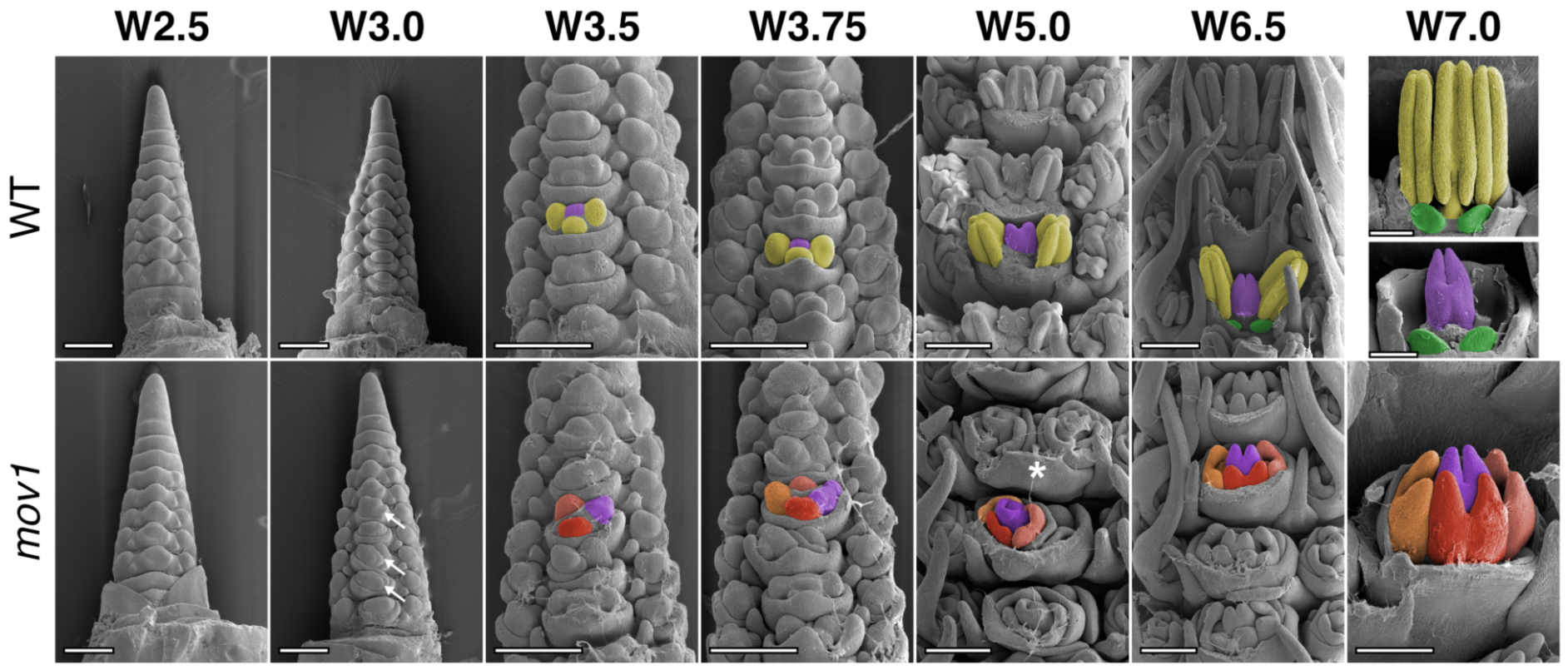
Scanning electron microscopy (SEM) of *mov1* inflorescence development. In wild-type (WT), primordia giving rise to stamens are false-coloured in yellow, cells giving rise to the carpel in purple and lodicules in green. In *mov1*, the central carpel (purple) is retained while the stamens are converted into additional carpels. Cells giving rise to the additional carpels are false-coloured in different shades of red/orange. White arrows in *mov1* (W3.0) indicate creases in the floral meristems, while white asterisk at W5.0 indicates separation of a single floral meristem into two distinct florets. Waddington stage is indicated for each developmental timepoint, from W5.0 lemma and/or stamens have been removed to expose the carpels. Scale bars: 200 µm.

To determine whether mutation of a B-class gene contributes to the *mov1* phenotype, a PCR-based strategy, consisting of PCR amplification followed by Sanger sequencing of the amplicon, was used to survey all B-class genes for presence/absence, structural and sequence variants in *mov1* plants relative to wild-type (cv. Steptoe). *HvMADS2* and *HvMADS4* did not show differences between genotypes when tested for amplicon size or sequence polymorphisms by PCR or by sequencing. In contrast, *HvMADS16* was not detected in *mov1* mutants (**Fig. S11A**). Genotyping by copy number analysis combined with phenotyping on a total of 583 plants demonstrated that *mov1* segregates as a single recessive Mendelian locus (3:1 - wild-type : multiovary) (**Table S3**). Absence of *HvMADS16* co-segregated perfectly with the *mov1* phenotype in all plants tested. These findings indicate that *mov1* lacks *HvMADS16*.

To define the size of the deletion surrounding *HvMADS16*, a PCR-based approach was used to test the presence/absence of neighbouring gene sequences based on annotations from the barley reference Morex genome Hv_IBSC_PGSB_v2. Overall, the *mov1* mutant appeared to be missing a region of approximately 0.95 Mb relative to wild-type (**Fig. S11C**). According to the reference sequence, this region is predicted to include three gene sequences: *HORVU7Hr1G091190* (40S ribosomal protein), *HORVU7Hr1G091200* (undescribed protein); and *HvMADS16* (**Table S4**). Consistent with a role of *HvMADS16* in inflorescence specification, transcripts were predominantly detected in developing wild-type inflorescences when tested by qRT-PCR across a Steptoe tissue series (**Fig. S11B**). On the other hand, transcripts for *HORVU7Hr1G091200* could not be detected in any of the tissues examined. Furthermore, a BLASTx query of the translated nucleotide sequence against NCBI non-redundant protein databases found no significant similarity to any protein in other species. Presence of gene *HORVU7Hr1G091190* encoding a 40S ribosomal protein could not be uniquely assayed due to the highly repetitive nature of its sequence. However, publicly available RNAseq data (Colmsee *et al*., 2015) indicate that *HORVU7Hr1G091190* is not expressed in several barley tissues including the inflorescence. Considering the specific homeotic conversion of floral organs in whorls 2 and 3, the absence of *HvMADS16* in *mov1* mutant plants and the role of B-class genes in other plant species, *HvMADS16* appears to be the most likely causal agent for *mov1*.

### HvSL1 and HvMADS16 differentially affect expression of floral regulators

To establish how the *HvSL1* (*mov2.g*) and *HvMADS16* (*mov1*) deletions might influence molecular pathways underlying stamen and carpel formation, qRT-PCR was performed on developing inflorescences from wild-type, *mov2.g* and *mov1* at stages W2.0 to W6.0 (**Fig. 8**), which encompass the specification and differentiation of the reproductive organs.

**Fig. 8.**
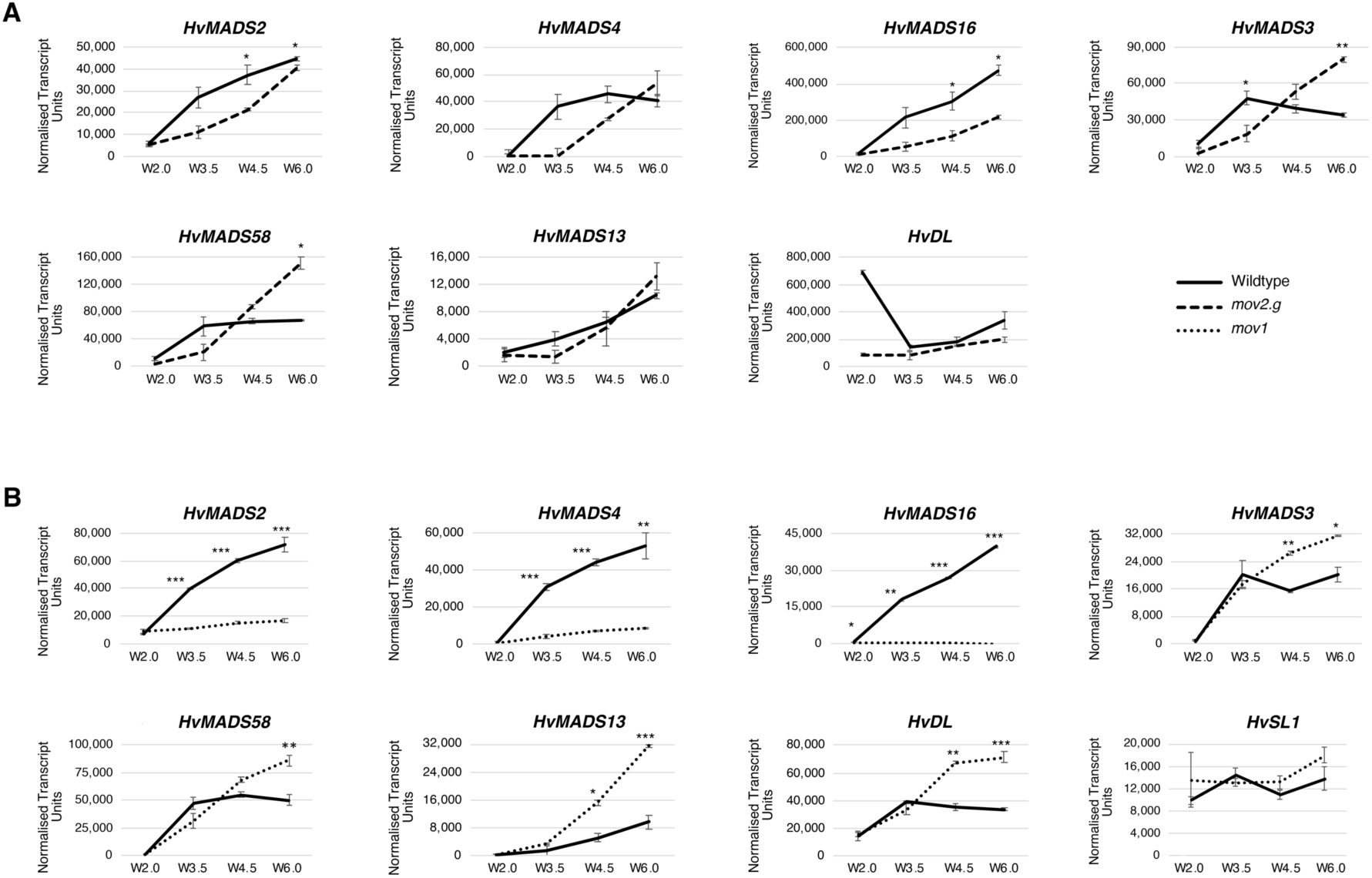
Transcript abundance of floral genes assessed by qRT-PCR in *mov2.g* and *mov1* developing inflorescences. Gene expression of B-class genes (*HvMADS2*, *HvMADS4*, *HvMADS16*), C-class genes (*HvMADS3*, *HvMADS58*), D-class gene *HvMADS13*, carpel gene *HvDL* and *HvSL1* was assayed in (**A**) *mov2.g* and (**B**) *mov1* inflorescences at stages W2.0 (double ridge), W3.5 (stamen primordia), W4.5 (carpel primordium) and W6.0 (stamen and carpel development). Error bars represent ± Standard Error. For each timepoint, two-tailed T-test P-values ≤0.05 (*), ≤0.005 (**) and ≤0.001 (***) are shown for differences between wild-type and *mov1*. For each sample n = 3 independent biological replicates.

When comparing transcript abundance for the three B-class genes in *mov2.g* (**Fig. 8A**), the most pronounced difference was observed for *HvMADS16*, which was significantly reduced in the mutant. However, despite an overall reduction of transcript abundance in *mov2.g*, expression still increased with development. A similar trend was observed for *HvMADS2*, while for *HvMADS4*, transcript abundance remained lower in *mov2.g* only until stamen primordia specification (stage W3.5). After stage W3.5, transcript abundance for *HvMADS4* steadily increased to match wild-type levels by stage W6.0. For C-class *HvMADS3* and *HvMADS58* MADS-box genes, transcript abundance in *mov2.g* showed a delayed accumulation relative to wild-type, followed by an increase eventually exceeding wild-type levels. Conversely, transcript abundance of ovule specific D-class gene *HvMADS13* and the barley orthologue of the rice carpel-specific YABBY transcription factor *HvDL* (*DROOPING LEAF*) remained largely unaffected in *mov2.g* samples.

In *mov1* (**Fig. 8B**), *HvSL1* expression did not significantly differ from wild-type. In contrast, expression of the B-class genes *HvMADS2* and *HvMADS4* was significantly lower in *mov1* from stage W3.5 onwards, and *HvMADS16* expression was absent. Conversely, expression of the C-class genes (*HvMADS3* and *HvMADS58*), D-class gene (*HvMADS13*) and the carpel-specific *HvDL* gene were all significantly increased in *mov1* compared to wild-type following stamen primordia specification (W3.5). To determine the spatio-temporal expression pattern during floret development, *in situ* hybridization of selected genes was performed in Steptoe and *mov1* mature inflorescences (**Fig. 9**). As expected, *HvMADS16* expression was undetectable in *mov1* inflorescences in any floral organ, whereas it was expressed in wild-type lodicules and stamens. The expression patterns of *HvMADS3* (C- class) and *HvMADS13* (D-class) genes were found to be very similar: in wild-type florets, expression was confined to the ovule, while in *mov1*, both *HvMADS3* and *HvMADS13* were expressed in multiple locations per floret. Notably, *HvDL* expression in wild-type was restricted to the carpels and to the abaxial side of the lemma but was absent in lodicules or stamens. In *mov1* florets, *HvDL* expression was more diffuse within the floret and remained in the lemma. For all genes tested, sense probes gave no observable signal (**Fig. S12**).

**Fig. 9.**
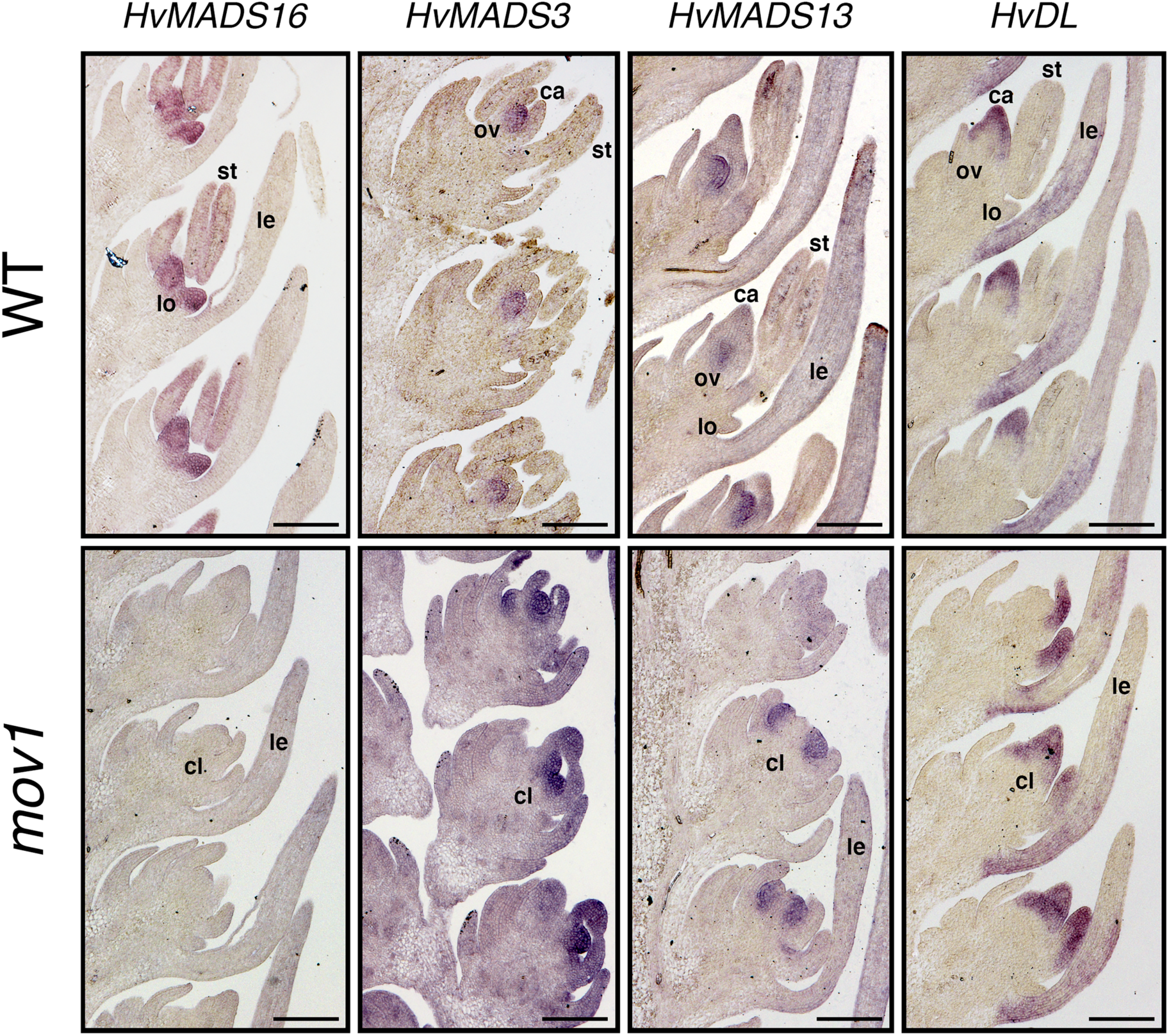
Spatial expression of floral homeotic genes in wild-type and *mov1* inflorescences. Expression patterns as detected by *in situ* hybridization are shown in wild- type (WT) and *mov1* inflorescences at stage W6.0 for genes *HvMADS16*, *HvMADS3*, *HvMADS13* and *HvDL*. Lemma (le), lodicule (lo), stamen (st), carpel (ca), ovule (ov) and carpel-like structure (cl). Scale bars: 200 µm.

## Discussion

### Deletion of the zinc-finger transcription factor HvSL1 underlies the mov2.g multiovary phenotype

The present study confirms the location of the *mov2* locus on the short-arm telomeric region of chromosome 3H (3HS) (Soule *et al*., 2000), and additionally refines the region to a 449 Kb interval in a *mov2.g* (cv. Steptoe) x Morex F_3_ population (**Fig. S3**). In contrast to previous reports, analysis of this region was unable to confirm suggestions that a MADS-box gene resides within the *mov2* locus. Instead, sequence analysis identified *HORVU3Hr1G003740*, referred to as *HvSL1*, as the most likely candidate for the *mov2.g* phenotype. Indeed, *HvSL1* is physically absent in *mov2.g* plants (**Fig. S6A**), a lesion that is consistent with fast-neutron mutagenesis (Kumawat *et al*., 2019). The absence of *HvSL1* completely co-segregates with the mutant phenotype and *HvSL1* expression is also completely abolished in *mov2.g* developing inflorescences (**Fig. S6B**). CRISPR-mediated *Hvsl1* knockout plants phenocopied *mov2.g*, providing additional evidence that the lack of *HvSL1* is causative for the *mov2.g* phenotype (**Fig. 4**).

*HvSL1* encodes a putative C2H2 zinc-finger transcription factor and is the likely orthologue of rice *STAMENLESS1* (*OsSL1*) (Xiao *et al*., 2009). Indeed, the *mov2.g* multiovary phenotype closely resembles that of *sl1* mutants in rice. However, notable phenotypic differences are present between *mov2.g* and *sl1*. Unlike rice *sl1*, a high rate of abnormal lodicules was not observed in *mov2.g* plants indicating some level of functional diversification between grass species. Functional and more pronounced differences are also identified between *HvSL1* and the respective orthologues in *Arabidopsis*, *JAGGED* (*JAG*) and *NUBBIN* (*NUB*). Although both *JAG* and *NUB* appear to be involved in correct stamen and carpel development, lack of function in these genes does not alter organ identity, but rather organ morphogenesis (Dinneny *et al*., 2006). Furthermore, the role of *JAG* and *NUB* as shape determinants is not restricted to the reproductive organs but also affects the outer floral whorls as well as vegetative tissues, indicating a more general role in proper lateral organ shape, especially for *JAG* (Ohno *et al*., 2004). This broad comparison indicates that barley *HvSL1* has evolved a function that more precisely controls specific floral organ identity in the *Triticeae* relative to both closely related monocots (rice) and the more diverse dicots (*Arabidopsis*).

The most prominent feature of *mov2.g* spikes is the ability to produce multiple seeds per floret upon cross-pollination, with up to three seeds developing simultaneously (**Fig. 1D, E**), a characteristic that has not been reported for rice *sl1*. Importantly, this indicates that most of the ectopic *mov2.g* carpels produce functional ovules and female gametophytes. Production of multiple seeds per floret is in accord with initial reports from the 1950s for multiovary *mo* mutants of the barley cv. Trebi (Moh and Nilan, 1953; Kamra and Nilan, 1959). However, unlike *mov2.g*, *mo* mutants exhibited incomplete penetrance of the multiovary phenotype, with some florets retaining one or two normal stamens, or showing incomplete conversion of stamens to carpels (Moh and Nilan, 1953). These phenotypic differences might be explained by allelic variation or distinct genetic backgrounds (Steptoe versus Trebi). Furthermore, florets of *mo* plants were commonly observed to contain a single lodicule (Kamra and Nilan, 1959), a characteristic that was not observed in *mov2.g* plants under our growing conditions. Overall, our results strongly suggest that *HvSL1* underlies the *mov2* locus and is essential for stamen development.

### Absence of HvMADS16 underlies the mov1 multiovary mutation

Previous mapping studies positioned the *mov1* locus on the centromeric region of chromosome 7H (Soule *et al*., 2000). The present work also identified a missing region of approximately 0.95 Mb on chromosome 7HL in *mov1* plants (**Fig. S11C**). However, the current study identified the *mov1* locus with higher resolution and significantly narrowed the critical interval to only three putative genes encoding: a 40S ribosomal protein, an undescribed protein (HORVU7Hr1G091200) and the HvMADS16 MADS-box transcription factor which shows high sequence identity (88.3%) to the rice B-class protein OsMADS16/SUPERWOMAN1 (**Fig. S11E**). Among the three missing genes in *mov1*, *HvMADS16* emerges as the most likely causative candidate. *mov1* plants show specific homeotic conversion of lodicules and stamens in whorls 2 and 3, respectively (**Fig. 6**), a phenotype that is consistent with the lack or reduced activity of B-class genes based on the ABC model of floral development. Additionally, the *mov1* phenotype is remarkably similar to B-class mutants such as *Arabidopsis apetala3* (*ap3*) (Jack *et al*., 1992), rice *superwoman1* (*spw1*) (Nagasawa *et al*., 2003) and maize (*Zea mays*) *silky1* (*si1*) (Ambrose *et al*., 2000), which all show transformation of petals/lodicules and stamens into bract-like and carpel-like organs and involve DEF-like genes. Furthermore, consistent with the observations by Soule *et al*., 2000, the 3:1 phenotypic segregation observed for individuals derived from *mov1* heterozygotes is typical for a single, recessive Mendelian locus, suggesting that no other loci are involved in the *mov1* phenotype (**Table S3**).

Additional molecular evidence supports our hypothesis that altered *HvMADS16* function underlies the *mov1* phenotype. Besides being physically absent in *mov1* (**Fig. S11A**), examination of *HvMADS16* transcript levels across different wild-type tissues found that the gene is preferentially expressed in inflorescences (**Fig. S11B**). At the tissue level, *HvMADS16* expression is specifically confined to lodicules and stamens as seen by *in situ* hybridization in wild-type (**Fig. 5C**). The very specific expression pattern is consistent with the observation that *mov1* only shows a major phenotype in floral development, but not in any other aspect of plant growth. The other gene found to be deleted in *mov1*, *HORVU7Hr1G091200*, is most likely a pseudogene as it is currently an unannotated low-confidence gene in the Hv_IBSC_PGSB_v2 Morex assembly, with no orthologues found within other species.

Another intriguing *mov1* phenotype is female sterility, since no seeds were obtained when *mov1* was used as female parent for crosses. Histological analysis suggested that all four carpels in *mov1* are non-functional because the ovules are unable to produce a fully differentiated embryo sac (**Fig. S10**). Instead, the ovule-like structures are filled with undifferentiated cell layers; this effect was also observed in wheat pistillody lines (Yamada *et al*., 2009) and in rice *spw1* (Nagasawa *et al*., 2003), suggestive of a potentially conserved role for HvMADS16 in ovule development. Interestingly, this contrasts with the *Arabidopsis* orthologue *ap3* mutant that produces viable seeds (Jack *et al*., 1992). It also contrasts with the lack of *HvMADS16* expression in ovules and suggests that the effect may be non-cell autonomous.

### Absence of HvSL1 and HvMADS16 disrupts the balance of the ABC model

Many studies have shown the vast and intricate interactions that exist among genes of the ABC model and the transcription factors driving floral development. Overall, the differences in gene expression we observed in *mov1* and *mov2.g* are consistent with the respective phenotypes.

As expected, *HvMADS16* expression in *mov1* is not detected by qRT-PCR and the signal from *in situ* hybridization is completely absent in the mutant inflorescences (**Figs 8B, 9**). Interestingly, the expression of the other B-class genes (*HvMADS2* and *HvMADS4*) in *mov1* is significantly reduced from a very early stage, from W2.0 (double ridge) onwards, perhaps indicating a tight regulatory relationship between these three genes in specifying lodicule and stamen identity (**Fig. 8B**). Additional evidence supporting this hypothesis comes from the overlapping expression of these genes in lodicule and stamen tissues as observed by *in situ* hybridization (**Fig. 5**). Through BiFC experiments we showed that HvMADS16 (DEF-like) is an obligate hetero-dimer with either GLO-like protein (HvMADS2 or HvMADS4) and that the GLO-like proteins do not interact with each other (**Fig. S8**). This observation is consistent with reports in numerous other species whereby DEF- and GLO-like hetero- dimers can activate their own expression and initiate a positive autoregulatory feedback loop (Schwarz-Sommer *et al*., 1992; Goto and Meyerowitz, 1994; McGonigle *et al*., 1996).

The reduced expression of all three B-class genes *HvMADS2*, *HvMADS4* and *HvMADS16* in *mov2.g* inflorescences is consistent with the lack of stamens (**Fig. 8A**). Nevertheless, it is worth noting that expression of these genes is not completely abolished and must be sufficient to drive normal ovule development in whorl 4, and lodicule development in whorl 2. Indeed, transcript abundance of both *HvMADS2* and *HvMADS4* reach levels that are comparable to wild-type by stage W6.0. Speculatively, the low levels of *HvMADS2* and *HvMADS4* at earlier stages could indicate a slower initiation of the positive autoregulatory feedback loop typical of B-class genes observed in other species, due to the low *HvMADS16* levels.

Overall, the expression of C-class genes (*HvMADS3* and *HvMADS58*), D-class genes (*HvMADS13*) and *HvDL* increases in *mov1* inflorescences (**Fig. 8B**). In wild-type, these gene classes are expressed in the innermost fourth whorl of the flower and function in specifying carpel (*HvMADS58, HvDL*) and ovule development (*HvMADS3, HvMADS13*). The transcriptional increase observed in *mov1* inflorescences can be explained by the combined effect of a minor repressive action of B-class genes on carpel and ovule-promoting genes, together with the expansion of the expression domain of these genes in the floral meristem (**Fig. 9**). The final effect of these expression abundance and domain changes is a floret with carpels in two consecutive whorls (whorl 3 and whorl 4).

The expression pattern of *HvDL* is consistent with the expression pattern of the rice orthologue *OsDL* (Nagasawa *et al*., 2003). In rice florets, *OsDL* expression has been shown to be confined to the carpel and to the central vein of the lemma (Ohmori *et al*., 2011). A very similar expression pattern was observed for *HvDL* in wild-type, with expression in the carpel and base of the lemma, thus suggesting a conserved function in carpel morphogenesis. In *mov1*, HvDL expression is more diffuse throughout the developing floret (**Fig. 9**).

Likewise, for *HvMADS13*, the spatial expression observed in wild-type inflorescences is consistent with the expression of the rice orthologue *OsMADS13* (Lopez-Dee *et al*., 1999). For both the barley ovule-specific genes, *HvMADS3* and *HvMADS13*, the increased expression detected in *mov1* inflorescences by qRT-PCR analysis may be explained by the additional ovules observed by histochemical analysis (**Fig. S10**). This corresponds well with the spatial pattern of expression of both genes in the mutant as visualised by *in situ* hybridisation, which shows distinct expression maxima throughout the floral meristem (**Fig. 9**). However, it is difficult to determine whether *MADS13*, *MADS3* and *HvDL* might be over-accumulating in their normal “domain” in *mov1* mutants. The expression of these ovule identity genes in ovule primordia is not sufficient to drive correct ovule formation, suggesting that their activity may be compromised, or that ovule fertility defects are manifested later during development.

In *mov2.g*, the increased abundance of carpel and ovule-specific transcription factors belonging to the C-class (HvMADS3 and HvMADS58) MADS-box genes (**Fig. 8A**) is consistent with the supernumerary carpels present in *mov2.g* compared to wild-type florets (**Fig. 1**). Despite the phenotype, expression levels of the D-class (HvMADS13) MADS-box gene and the carpel-specific HvDL gene remain unaffected in *mov2.g* with respect to wild-type (**Fig. 8A**). This contrasts with *mov1*, and confirms that an increase in ovule number alone does not always relate to increased *HvMADS13* and *HvDL* expression. These two genes may play a minor role compared to other genes in the formation of additional ovules and carpels in *mov2.g*. Alternatively, HvDL in *mov2.g* may have slower expression dynamics. Indeed, most of the investigated genes seem to have a delayed response in *mov2.g* compared to wild-type.

### HvSL1 and HvMADS16 functionally overlap during floral specification

As seen by SEM analysis, phenotypic differences in *mov2* and *mov1* inflorescences appear very early in flower development (**Figs. 2, 7**). This is consistent with the spatial expression pattern of both *HvSL1* and *HvMADS16* transcripts in wild-type inflorescences as observed by *in situ* hybridization whereby signals are detected in young meristems before the differentiation of floral organs (**Figs. 3, 5**). It is also in accordance with the temporal expression pattern as shown by qRT-PCR, as expression of both *HvSL1* and *HvMADS16* in a wild-type inflorescence is first detected at double ridge (stage W2.0) and gradually increases until differentiation of the reproductive organs (W6.0) (**Figs S6B, 8**). When considering the arrangement of floral organs into whorls, *mov2.g* plants show distinct homeotic conversion of whorl 3 organs (stamens) into supernumerary carpels, while floral organs in the other whorls remain unaffected (**Fig. 1**). For this reason, we propose that *HvSL1* is specifically necessary for correct stamen development.

Studies in rice have suggested that OsSL1 may act as a positive upstream regulator of *OsMADS16* transcription (Xiao *et al*., 2009). Considering that *HvSL1* expression temporally precedes *HvMADS16* and given the overlapping expression domains within stamen primordia, we wanted to confirm whether *HvSL1* is involved in the correct regulation of B-class genes, particularly *HvMADS16* in barley. Indeed, the presence of the HvSL1 transcription factor appears to positively influence endogenous *HvMADS16* expression in transfected protoplasts (**Fig. S9C**). Consistent with the hypothesis that *HvSL1* acts upstream of *HvMADS16*, expression of *HvMADS16* is downregulated in *mov2.g* even before stamen primordia initiation, whereas the expression profile of *HvSL1* does not significantly differ from wild-type in *mov1* plants (**Fig. 8**). These results suggest a conserved regulatory network with rice in controlling stamen identity.

### A model for barley stamen specification

Although the ABC model was initially proposed as a universal model to explain flower development, differences have been found across plant species, even among members belonging to the same family as is the case for rice and barley (*Poaceae*). For example, an expression atlas through inflorescence development of 34 barley MIKc MADS genes highlighted how expression of some genes strongly deviates from predicted patterns, such as the apparent absence of antagonism between A- and C-class genes in barley (Kuijer *et al*., 2021). Moreover, some classes have undergone gene duplications acquiring grass-specific functions as seen for rice A-class genes which specify palea and lodicules (Callens *et al*., 2018), while other classes underwent partial sub-functionalization following duplication, as is the case of the rice C-class genes *OsMADS3* and *OsMADS58* (Kang *et al*., 1995; Yamaguchi *et al*., 2006, Kuijer *et al*., 2021). Indeed, in the current study, the spatial expression pattern of *HvMADS3* detected by *in situ* hybridization also highlights a potential sub-functionalization of C-class genes in barley. Both the rice (*OsMADS3*) and maize (*ZMM2*) orthologues of *HvMADS3* have been reported to be strongly expressed in stamen primordia (Dreni *et al*., 2011) and anthers (Mena *et al*., 1996), respectively. Furthermore, it has been shown that *OsMADS3* plays an important role in specifying stamen identity, as well as in repressing lodicule formation (Yamaguchi *et al*., 2006). In contrast, in wild-type barley inflorescences *HvMADS3* expression was specific to the ovule primordia (**Fig. 9**). Furthermore, *HvMADS3* expression was shown by qRT-PCR to increase in *mov1* inflorescences (**Fig. 8B**), which wouldn’t be expected if the function of *HvMADS3* were conserved with respect to its rice and maize orthologues.

Other genes in barley have adopted different roles in addition to floral specification. This is the case for the A-class gene *HvMADS14* which controls vernalization-induced flowering, acting as the *VERNALIZATION1* (*VRN1*) equivalent in other temperate cereals (Trevaskis *et al*., 2003). Another example can be found in *HvMADS1* (E-class), which has been recently reported to direct plant thermomorphogenesis by acting on maintaining unbranched spike architecture at high temperatures (Li *et al*., 2021). Conversely, B-class function, particularly stamen development, appears to be extremely conserved among domesticated grasses such as rice, maize and barley, and may be independent of any environmental adaptation to temperate or tropical climates. The effect on stamens observed in *HvMADS16* closely reflects what has been reported for rice *spw1* (Nagasawa *et al*., 2003), maize *si1* (Ambrose *et al*., 2000) and more broadly *Arabidopsis ap3* (Jack *et al*., 1992).

Based on the results obtained in this study, together with previous reports in barley and other plant species, we formulate a testable model for barley stamen specification (**Fig. 10**). In this model, HvSL1 acts predominantly in whorl 3 and promotes *HvMADS16* in wild-type (**Fig. 10A**). The B-class genes *HvMADS16*, *HvMADS2* and *HvMADS4* are expressed in floral whorls 2 and 3 and form DEF-GLO hetero-dimers. Based on the floral quartet model, the hetero-dimers associate in higher order protein complexes with A- and E-class genes leading to the formation of lodicules in whorl 2. In whorl 3, instead, the floral quartet complex forms between B-, C- and E-class genes promoting stamen development while concomitantly repressing carpel formation. The lack of *HvSL1* in *mov2.g* plants results in a shift in this balance (**Fig. 10B**). At a molecular level, we speculate that B-class activity in *mov2.g* flowers is predominantly maintained in whorl 2 where enough HvMADS16 is present to develop normal lodicules. At the same time, the reduced levels of HvMADS16 in whorl 3 are insufficient to successfully form functional B-class hetero-dimers. Consequently, the predominant quaternary complexes forming in whorl 3 are between C- and E- class genes, which promote carpel formation. Concomitant expansion of ovule-promoting genes such as *HvMADS3* to whorl 3 may then explain multiple carpels having a functional female gametophyte.

**Fig. 10.**
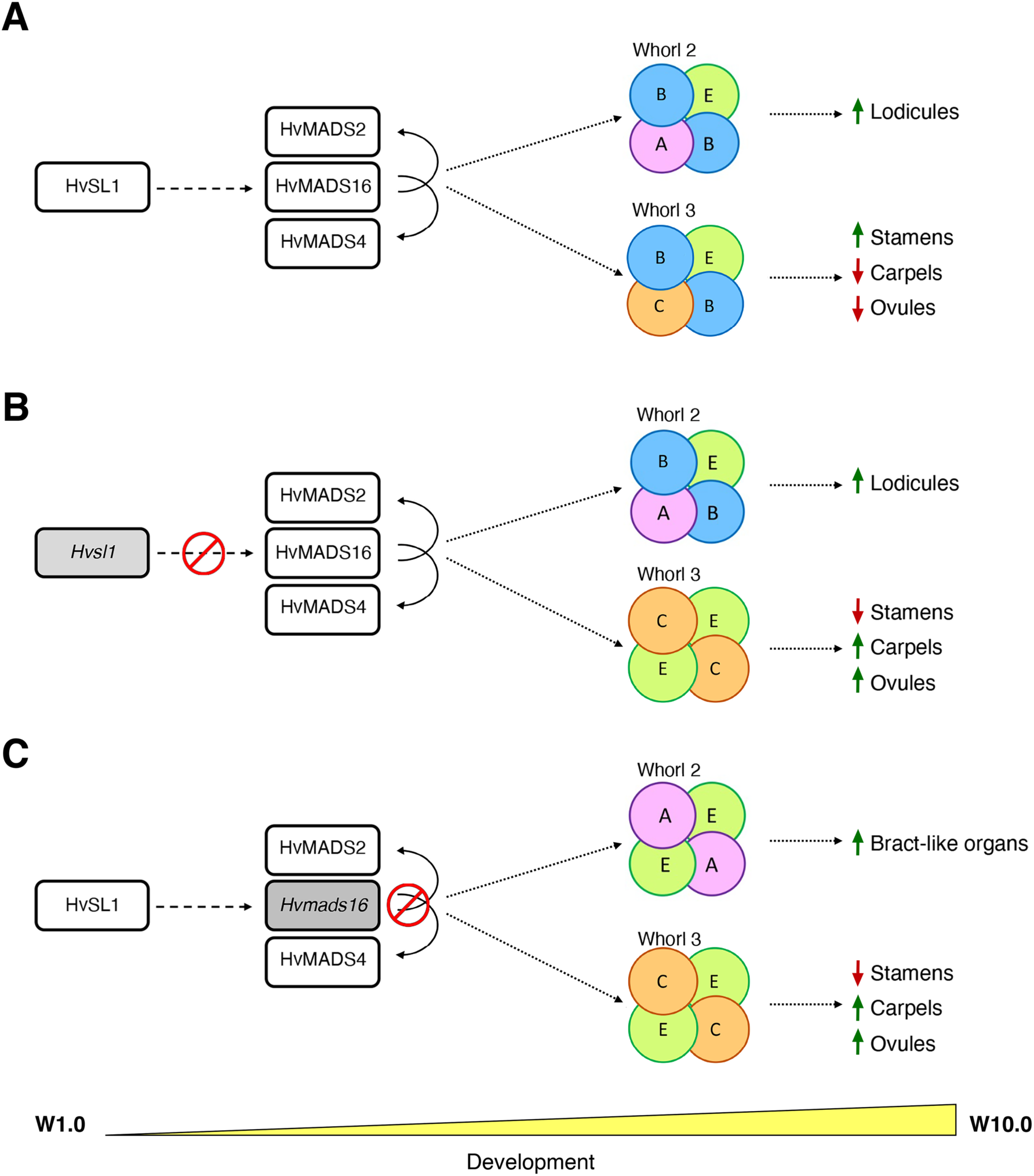
Model for barley stamen specification. (**A**) As inflorescence development progresses in wild-type, genes of the ABC model combine in floral quartets to specify lodicules in whorl 2 and stamens in whorl 3, with putative repression of carpel and ovule- specific genes. (**B**) The absence of *HvSL1* in *mov2.g* plants and (**C**) of *HvMADS16* in *mov1* plants affects the balance and composition of the floral quartets that form in these whorls, leading to an altered specification of the resulting floral organs. Dashed arrows indicate direct or indirect interaction, solid arrows indicate direct interactions found in this study, while dotted arrows indicate a process or consequence. For each mutant, the missing protein is indicated in grey and the red symbol indicates the affected interaction.

In *mov1*, *HvMADS16* is absent. The remaining B-class GLO-like proteins HvMADS2 and HvMADS4 do not interact and therefore cannot compensate for the absence of HvMADS16. Thus, the only protein complexes that can form in whorl 2 are between A- and E-class genes, and between C- and E-class genes in whorl 3. As a result, bract-like organs develop in whorl 2 and carpel formation is promoted in whorl 3, leading to the multiovary phenotype observed in *mov1* (**Fig. 10C**).

In conclusion, we have reported the fine characterization of two multiovary loci, *mov2* and *mov1*, and identified likely causative genes, namely a C2H2 zinc-finger transcription factor termed here *HvSL1*, and the B-class gene *HvMADS16*. Although both mutants exhibit a similar multiovary phenotype, the difference in fertility is remarkable: while *mov1* is completely sterile, *mov2* can produce multiple seeds per floret. This indicates that the presence of multiovary is not necessarily indicative of fully functional reproductive development in barley. Studying the molecular changes underlying these differences will improve our understanding of floral development at a species-specific level, and expand the tools available for modification of cereal flowers for breeding.

## Supplementary data

Supplementary data are available at *JXB* online.

**Table S1A.** Primer sequences and Taqman probes used for copy number analysis to genotype *mov2.g* and *mov1* plants.

**Table S1B.** Sequence of KASP^TM^ markers on chromosomes 3H used to map the *mov2* locus.

**Table S1C.** PCR primer sequences for testing the presence of genes upstream and downstream from *HvSL1* on chromosome 3H.

**Table S1D.** PCR primer sequences for testing the presence of barley B-class genes.

**Table S1E.** PCR primer sequences for testing the presence of genes upstream and downstream from *HvMADS16* on chromosome 7H.

**Table S1F.** qRT-PCR primer sequences.

**Table S1G.** Primer sequences for *HvSL1* CRISPR knockout. gRNA sequence is underlined.

**Table S1H.** Primer sequence for cloning of *in situ* hybridization antisense and sense probes.

**Table S1I.** Primer sequences for BiFC cloning.

**Table S2.** Annotated genes present in the mapped *mov2* critical interval between flanking markers chr3H_9748112 and chr3H_10289104.

**Table S3.** Observed segregation ratios of *mov1* phenotype in heterozygote growing material.

**Table S4.** Genes on chromosome 7H tested by PCR.

**Fig. S1.** Histological sections of *mov2.g* carpels.

**Fig. S2.** Spike morphology of wild-type and *mov2.g* plants.

**Fig. S3.** Mapping of the *mov2* locus in a *mov2.g* x Morex bi-parental population.

**Fig. S4.** Heatmap of gene expression for genes in the critical *mov2* interval.

**Fig. S5.** *HvSL1* and *OsSL1* gene models and alignment.

**Fig. S6.** *HvSL1* deletion in *mov2.g*.

**Fig. S7.** CRISPR design and analysis of *Hvsl1*-knockout plants.

**Fig. S8.** BiFC assays showing interaction between barley B-class genes.

**Fig. S9.** Transfection efficiency and transcript abundance in barley protoplasts.

**Fig. S10.** Histological sections of *mov1* carpels.

**Fig. S11.** Characterization of *mov1* deletion and *HvMADS16*.

**Fig. S12.** *In situ* hybridization with sense probes on wild-type and *mov1* inflorescences.

## Acknowledgements

This research was supported by funding from the Australian Research Council (M.R.T.; DP210103491), the Australian Centre for Plant Functional Genomics (R.W. & U.B.; ACPFG), the University of Adelaide and the Australian Government who provided a Research Training Program Stipend Ph.D. scholarship to C.S. The authors are grateful to Dr Margaret Pallotta and Mr Chao Ma for technical advice and to the Adelaide Microscopy staff at the University of Adelaide.

## Author contributions

C.S., R.W., U.B. and M.R.T. conceived the research. R.W., U.B. and M.R.T. supervised the project. C.S. carried out the experiments. N.J.S. performed the qRT-PCR and assisted in identifying KASP^TM^ markers for mapping. X.Y. assisted with *in situ* hybridization and technical advice on cloning. C.S. wrote the manuscript; all authors edited and reviewed the manuscript.

## Data availability

All data supporting the findings of this study can be found within the paper and its supplementary materials published online. They are also available from the corresponding author, A/Prof. Matthew R. Tucker, upon request.

